# The role of hub neurons in modulating cortical dynamics

**DOI:** 10.1101/2021.06.11.448003

**Authors:** Eyal Gal, Oren Amsalem, Alon Schindel, Michael London, Felix Schürmann, Henry Markram, Idan Segev

## Abstract

Many neurodegenerative diseases are associated with the death of specific neuron types in particular brain regions. What makes the death of specific neuron types particularly harmful for the integrity and dynamics of the respective network is not well understood. To start addressing this question we used the most up-to-date biologically-realistic dense neocortical microcircuit (NMC) of rodent, which has reconstructed a volume of 0.3 mm^3^ and containing 31,000 neurons, 36 million synapses, and 55 morphological cell types arranged in 6 cortical layers. Using modern network science tools, we identified “hub-neurons” in the NMC, that are connected synaptically to a large number of their neighbors and systematically examined the impact of abolishing these cells. In general, the structural integrity of the network is robust to cells’ attack; yet, attacking hub neurons strongly impacted the “small worldness” topology of the network, whereas similar attacks on random neurons have a negligible effect. Such hub-specific attacks are also impactful on the network dynamics, both when the network is at its spontaneous synchronous state and when it was presented with synchronized thalamo-cortical visual-like input. We found that attacking layer 5 hub neurons are most harmful to the structural and functional integrity of the NMC. The significance of our results for understanding the role of specific neuron types and cortical layers for disease manifestation is discussed.

## Introduction

Research at the macro- and meso- scale brain anatomy has demonstrated a clear connection between structure-to-function. Indeed, the global network structure of the brain was shown to be altered in diseases such as schizophrenia, (Rubinov and Bullmore, 2013), and bipolar disorder (Syan et al., 2018) and other (Stam, 2014). Yet, pathology takes place at the microscale, at the cellular and synaptic level architecture of neuronal microcircuits. How does the connectomics at this level shape the dynamics and functionality of biological circuits is indeed a key question in neuroscience (Abbott et al., 2020; Turner et al., 2020). Of particular interest is the impact of structural disruption of the connectome, whether due to natural aging or due to diseases. These two types of disruptions are rather different. Whereas a nonselective general reduction in the number of cells was found in the aging brain, recent studies showed selective cell vulnerability associated with certain pathologies. For example, a significant decrease in the number of specific cell types in cortical areas in Alzheimer’s disease (Stranahan and Mattson, 2010; Fu et al., 2018; Murray et al., 2018), multiple sclerosis (Schirmer et al., 2019) and Parkinson (Hammond et al., 2007).

Experimental investigation of the role of specific cell populations in the neocortex has advanced significantly in recent years. Optogenetic methods (Deisseroth, 2015) together with genetic dissection of specific neurons (Luo et al., 2018) enables precise recording and manipulation (silencing and activating) of specific neuronal populations. Manipulating the neural activity of specific cell types during *in vivo* experiments is presently used to manipulate animal behavior (Guo et al., 2015; Carrillo-Reid et al., 2019; Robinson et al., 2020), but the effects of such cell-type-specific manipulations on circuit dynamics are rarely characterized at the network scale (Cardin et al., 2009; Pouille et al., 2009; Adesnik and Scanziani, 2010; Xue et al., 2014; Bitzenhofer et al., 2017). Consequently, we currently lack understanding of the role of particular cell populations, e.g., following specific diseases, in shaping neural network dynamics and eventually network functionality.

To address this gap, we hereby utilized theoretical approaches to explore the correlates between microcircuitry structure and function. Towards this end we simulated the most up-to-date biologically-realistic dense digital reconstruction of a neocortical microcircuit, NMC (Markram et al., 2015). This 0.3 mm^3^ cortical circuit contains some 31,000 neurons, 36 million excitatory and inhibitory synapses, and 55 morphological cell types (m-types). This model circuit enables an unprecedented opportunity to directly investigate the impact of network structure on system dynamics by introducing cell-specific and layer-specific attack/damage while measuring the collective neural activity under different physiological conditions.

Of particular interest is the question of whether certain cell types (“hub cells”) that either receive or make a more-than-average number of synapses are particularly impactful for network dynamics and, perhaps also for neurological diseases. Our recent experimentally-based theoretical work has identified such “hub neurons” in the NMC (Gal et al., 2017). It was shown that, among the ~55 m-types that constitute the mouse somatosensory cortex, only a limited number of cell types may consist of rare hub neurons having a significantly high number of out-going and in-coming connections, e.g., the thick tufted L5 pyramidal cells. The existence of such cell-type-specific wiring specificities was found essentially inevitable (Gal et al., 2019) and highly connected cells were shown to have a dominant role in shaping the cortical circuit dynamics (Setareh et al., 2017; Luccioli et al., 2018). Is the death (“attack”) of these cells in realistic cortical microcircuits more harmful to the network dynamics as compared to that of other cell types?

Using network science tools, we first analyzed how the circuit topology was impacted following a cell attack. This analysis includes computing the mean shortest path between two nodes in a network before and after such attacks; this measure is related to information flow in the network. Another respective measure of the network topology is its Small-World characteristic. Reduction in the “small-worldness” of the networks might imply a reduction in efficiency of information exchange and capacity for associative memory (Bullmore and Sporns, 2009). The impact of cell attack on additional topological measures is also examined in this study.

We next used several measures to evaluate network dynamics such as mean firing rate, coefficient of variance (CV), SPIKE-synchronization (Kreuz et al., 2015), etc. Each measure provides us with different aspects of the circuit activity and, hence, helps us understand how targeted attacks on hub neurons are more disruptive to the network functionality than attacking the same number of neurons randomly. Combining these results with structural network measures following cells’ attack shed new light on the robustness of the neocortical microcircuit connectivity to a variety of attacks and, at the same time, on the functional sensitivity of the network to some of these attacks. These findings provided important insights into the impact of the death of specific cell types (e.g., due to certain diseases) on the dynamics and functionality of local cortical microcircuits.

## Results

### Hub neuron attacks impact the NMC small-world topology

To explore the structural and functional impact of attacking highly connected “hub-neurons” we started by ranking all neurons according to their total degree (total number of pre- and post-synaptic cells connected to a given neuron). We then removed (attacked) different quantities of these neurons, starting from the highest degree hub cells to the lowest degree.

To quantify the structural effect, we first measured the overall connectedness of the network, as captured by the size of the network’s largest connected component (giant component size; see **Methods**). We highlight two extreme outcomes on network architecture following hubs attack (**Fig. 1A**). On one extreme, the removal of hub neurons may completely break the network into multiple smaller unconnected components (**Fig. 1A** top; “breakable”). In the other extreme, the removal of hub neurons will not break the network and the remaining neurons will remain in one connected giant component. (**Fig. 1A** bottom; “unbreakable”). A finer structural feature utilized here, which relates to the efficiency of network communication and computation, is the “small-world topology” of the network. This measure relies on two opposing requirements: a short path length between any pair of nodes/neurons (**Fig. 1B**, right) and clustered interconnectivity, *c*, within groups of nodes (**Fig. 1B**, left), see (Watts and Strogatz, 1998). Thus, the “small-worldness” of a neural network reflects the degree in which it balances the needs for global integration and local segregation of neural information (Sporns, 2013a).

**Figure 1.**
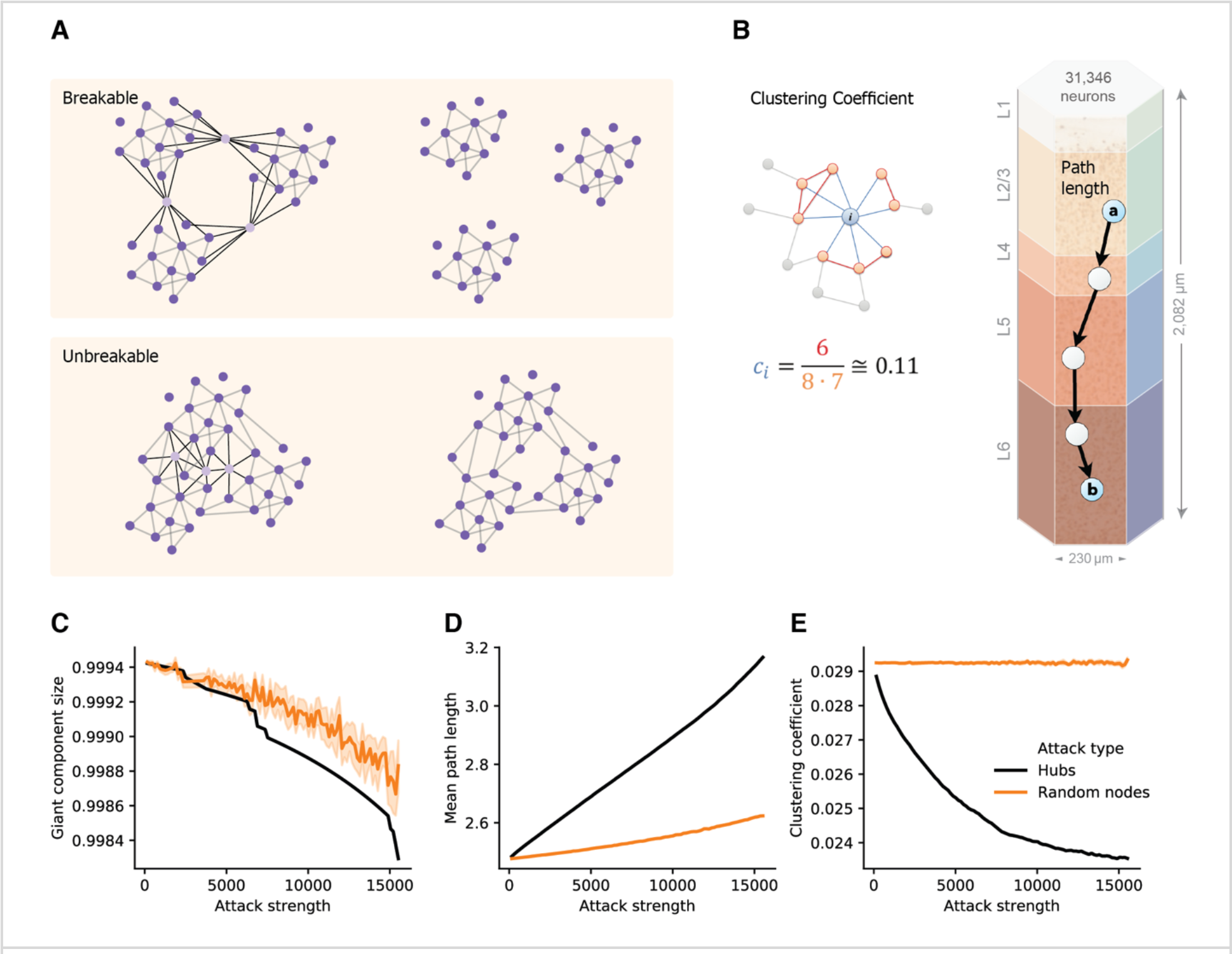
Structural disruption of cortical microcircuits following attacks on hub-neurons. **(A)** Schematic illustration of two network architectures having different sensitivity to a similar targeted hubs attack. **Top**: the case of breakable network whereby an attack on 3 hub neurons disintegrates the network into 3 separate sub-networks. **Bottom**: the case of an unbreakable network which, after an attack on 3 hub neurons, the network remains fully connected. **(B)** Schematic illustration of the two features used to define “small world” networks. **Right.** The shortest path length connecting two nodes (cells “a” and “b”, overplayed on the modeled neocortical microcircuit). **Left.** The local clustering coefficient of a node *i*, *c*_*i*_, which is the density of connections among the neighbors of this node. In the example shown, node i, has 8 neighbors; among them only 6 connections out of all 8·(8-1) = 56 possible connections (*c_i_* = 6/56 ≅ 0.11). **(C)** The size of the giant component in the NMC as a function of the number of hubs attacked (Attack strength), black line, compared to the corresponding random attacks (orange line), demonstrating that the NMC network is “unbreakable”. **(D)** Mean path-length and (**E**) mean clustering coefficient following hub attacks (black) versus random attack (orange). These two features are particularly sensitive to hub attack. Light orange depicts 95% confidence interval in all figures.

We found that the giant component of the NMC is not broken by hub-attack as its size only negligibly decreased when a large number of hub-neurons are attacked (**Fig. 1C**, black line), showing that the global integrity of the circuit is robust to such attacks. Moreover, the effect was similar to respective random cell-attacks (**Fig. 1C**, orange line). This implies that the NMC circuit is of the unbreakable type. In contrast, the small-world topology of the circuit is more sensitive to attacks on hub neurons. For example, attacking 5,000 hub neurons increased the path length from 2.48 to 2.69 (8% increase, **Fig. 1D**) while reducing *c* from 0.029 to 0.025 (12% reduction, **Fig. 1E**). To test whether this disrupted small-world topology is expected by chance and resulting merely due to the number of eliminated nodes, we performed control random attacks with matching number of nodes (**Fig. 1C-E**). We found that the disruption of both path-length and clustering due to hub attacks were significantly stronger than that expected from the random attacks (p < 0.001 for both, two-tailed Wilcoxon rank sum test; n_1_ = 10, n_2_ = 100; **Methods**). Additionally, the observed disruptions were found significant (P < 0.001, two-tailed Wilcoxon rank sum test; n_1_ = 10, n_2_ = 100; Methods) compared to that of randomly attacked networks with similar numbers of eliminated edges (**Supplementary Figure S1**).

### Hub neurons are key for network synchrony

The structural analysis has uncovered the disruptive effect of hub attacks on the small worldness properties of the neocortical microcircuit. However, the functional implications of such structural changes are not trivial. To elucidate the functional impact of hub neurons on network dynamics, we simulated MNC networks following different cells’ attacks and examined various functional features of network activity.

It has been shown (Markram et al., 2015) that at an extracellular calcium concentration of 1.4 mM the NMC network generates spontaneously synchronous bursts at ~1 Hz (**Fig. 2A** and see **Methods**). At this synchronous state, the cells’ mean firing rate is 3.7 Hz, their CV is 2.16 and SPIKE-synchronization measure is 0.25 (see **Methods**). We simulated the circuit after removing 2,977 hub neurons (**Fig. 2B**) or 2,977 random neurons (**Fig. 2D**). Removal of hub neurons markedly reduced the number of bursts in the network (**Fig. 2B**), whereas the bursting properties of the network were unaltered for the respective random attack (**Fig. 2D**). Increasing further the number of attacked hub-neurons to 7,993 completely abolished the bursting activity of the network, shifting it to the asynchronous state (**Fig. 2C**) whereas it did not change the burstiness in the case of random cells removal (**Fig. 2E**). **Figure 2F** summarizes the change in burst number due to different strengths of attacks. In addition to the reduction in burst activity due to removal of hub cells from the circuit, the correlation of variation (std divided by mean ISI), CV, firing rate and spike-synchronization all showed a stronger reduction compared to removal of random neurons (**Fig. 2G-I**).

**Figure 2.**
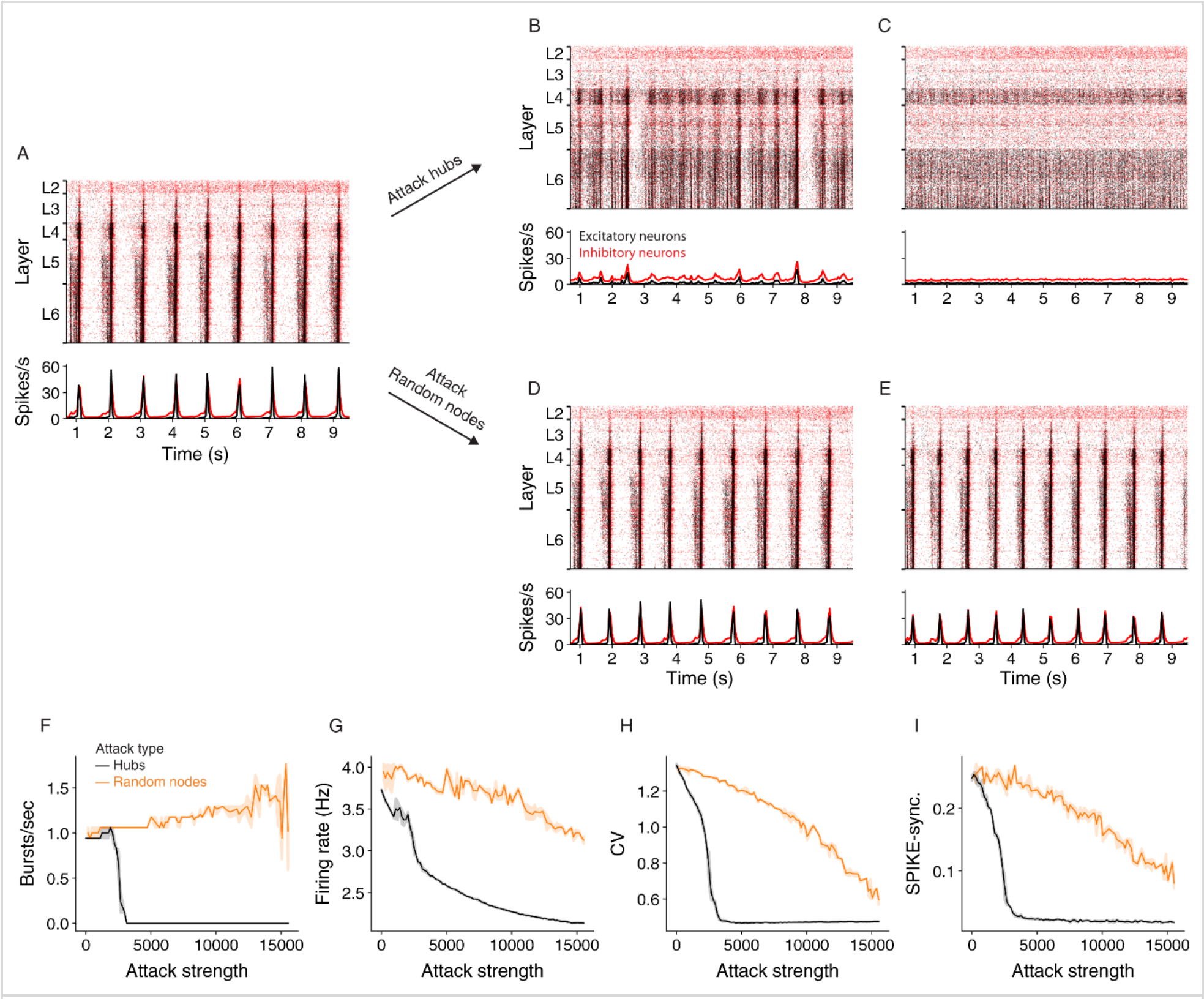
Effect of hub attacks on the dynamics of the NMC network in the synchronous regime. **(A)** Raster plot (top) and time histogram (bottom) of the NMC spiking activity during spontaneous bursting state (**Methods**). In this state, all cortical layers tend to burst synchronously at about 1Hz, with lower layers starting to fire earlier. **(B,C)** Same as (A) after attacking 2,977 and 7,993 hub neurons, respectively. **(D,E)** As in **(B,C)** but for respective random attacks. **(F-G)** Impact of hubs attack (black) versus random attack (orange) on network activity as a function of attack strength. **(F)** Impact on the number of bursts/sec. **(G)** Impact on average firing rate. **(H)** The impact on correlation of variation. **(I)** On global SPIKE synchronization measure. For all measures, hub attack is significantly more impactful. (light orange depicts 95% confidence interval).

Because hub neurons are mostly excitatory (Gal et al., 2017), hub attacks primarily remove excitatory neurons from the network. Indeed, in all analyzed hub attacks, ranging up to 15,000 neurons, the percentage of excitatory neurons in the attacked neurons was above 97%, but when attacking random nodes, we converge to the full circuit distribution of E/I neurons (85% excitatory). To address this discrepancy, we performed random attacks that matched both the number and the E/I identity of nodes (**Supplementary Fig. 2** and **Methods**). Indeed, this attack was more disruptive than the completely random attacks, but still less than the hub attacks. We also show that the number of edges attacked is not the main factor for this effect (**Supplementary Fig. 2**). This analysis demonstrates that, in the synchronous state, the impact of a neuron on the generations of collective synchrony in the NMC is more affected by their embedding in the network (“hubiness”) rather than by their physiological effect (the network E/I distribution).

Finally, we repeated the above analysis also for the asynchronous state, which is induced by setting the calcium concentration in the MNC simulations to 1.25mM (**Methods** and Figure 15 in Markram et al., 2015). At this state, the difference between attacking hubs versus random neurons is still significant, but not prominent as in the asynchronous case (**Supplementary Fig. 3; Methods).** Indeed, the asynchronous state is, in general, more robust to cell-attacks.

### Functional implication of layer-specific hub attacks

We showed that hub neurons are more effective in driving network synchrony. These hubs potentially belong to multiple cell-types at the different layers. To further detail the impact layer-specific hubs, we measured the connectivity among excitatory and inhibitory cells within and between layers (**Fig. 3A**). To compactly examine the connectivity among all 55 cell types in the NMC, we employed a force-directed graph drawing algorithm, whose 55 nodes depict the cell types whereas edge strengths correspond to the pairwise connection probability. In this presentation, tightly connected nodes will tend to appear closer. Inspecting the original network, the existence of large cell-type groups and clusters can be seen within each layer (**Fig. 3B** large nodes). After attacking 15,000 hub neurons, several changes were prominent (**Fig. 3C**). The network layout had spread more widely, indicating that the strength of the connections between cell types is reduced. Additionally, the large nodes of L5 almost disappeared, hinting to their possible impact on disruption of network functionality (see below). The interactive version of this algorithm implementation on the NMC is available online for further examination at intermediate levels of hub attacks.

**Figure 3.**
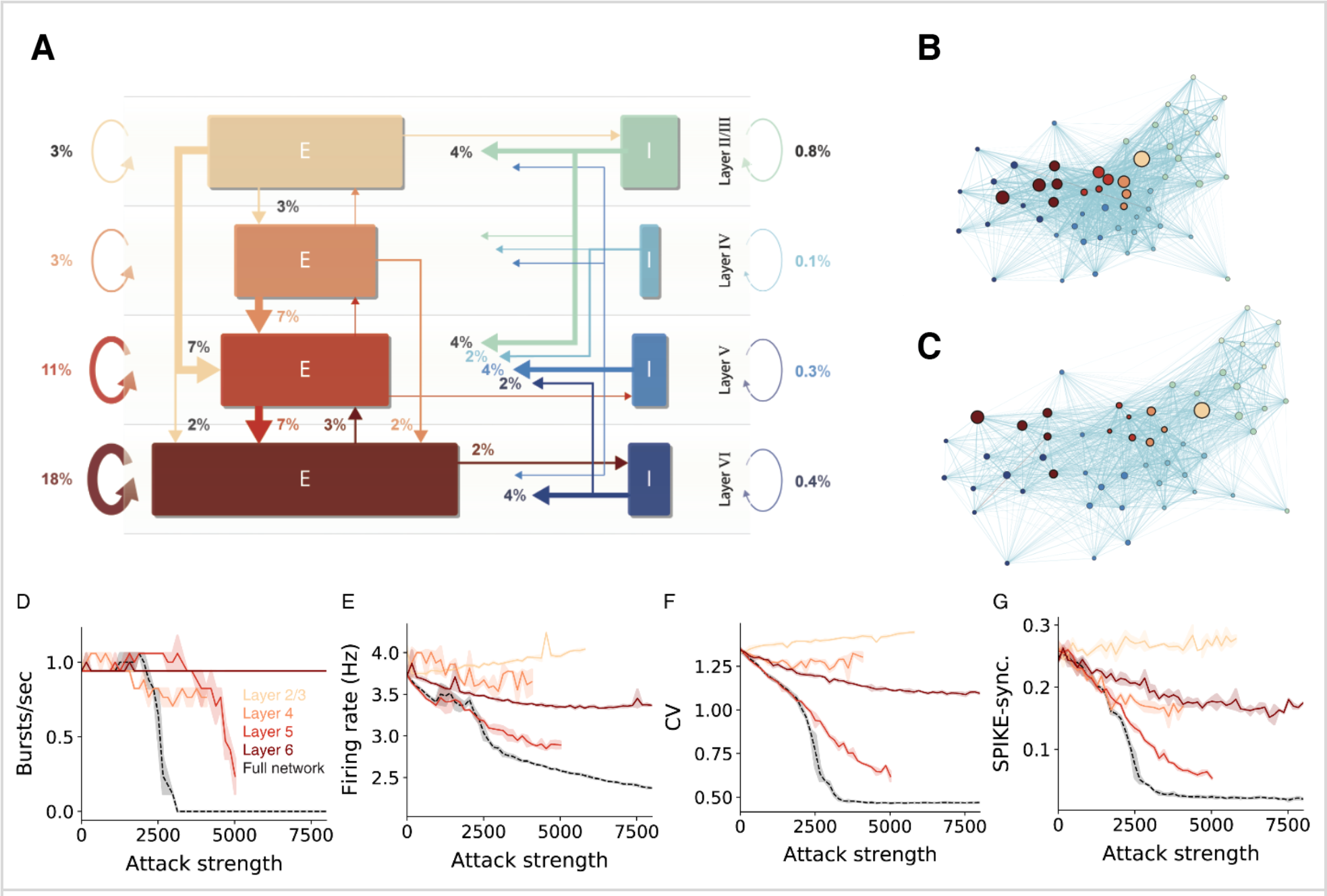
Sensitivity of network dynamic to hub attack is mostly attributed to attacks on L5 PCs. **(A)** Percentages of synapses for excitatory (brown-red arrows) and inhibitory (blue-green arrows) connections in the NMC microcircuit (layer 1 omitted). Arrow width and corresponding numbers indicate the percentage of total synapses formed by this pathway (omitted for pathways with <1% of synapses). The total percentage of plotted synapses is 98%; the remaining 2% originate in layer 1). Rectangle sizes are proportional to the sizes of the corresponding number of excitatory or inhibitory populations. **(B)** The connectivity among layers and cell-types is visualized using the force-directed graph algorithm (**Methods**). The network is composed of 55 morphological cell-types (circular nodes, colors match to that in A); edges strength corresponds to pairwise connection probabilities. Strongly connected cell-types are displayed closer in space. **(C)** Same as B, but for the network following an attack on 15,000 hub neurons. Note the disappearance of the large nodes in Layer 5. **(D-G)** Different quantification of network activity for layer-specific hub attack, global hub attack and random nodes attack. **(D)** Number of bursts/sec, **(E)** Average network firing rate, **(F)** Correlation of variation, **(G)** On global synchronization measure (**Methods**). Note that attack of L5 pyramidal cells is the most disruptive layer attack.

To test the importance of the different layers for network activity we performed layer-specific hub attacks by simulating the circuit while removing hub neurons in specific layers. We found that, in general, attacking L5 hub-neurons is the most distributive attack as it caused the largest change in all the functional measures used (**Fig. 3D, F-G,** but see the impact of L4 in **3E**). When revisiting Figure 3A one sees that L5 excitatory cells are the most interconnected population in the circuit (the overall percentage of incoming and outgoing connections); this is probably the reason for the high functional influence of L5 attack. L6 attacks also resulted in a large change of the functional measurements, although it did not influence the number of bursts.

We summarize this section by noting that attacking hub neurons is most disruptive to network dynamics when hub neurons are attacked at all layers (**Fig. 3D-G**, dashed black line). Layer 5 is the most-sensitive layer to such an attack. As neurons in different cortical layers belong to different genetic types (Gouwens et al., 2019; Yuste et al., 2020), this result shows that, although the same number of neurons might be degraded due to different pathologies that target specific genetic cell types (e.g., L5 thick-tufted pyramidal cells), they will have a very different impact on the overall dynamics of the cortical network and, thus, on the manifestation of specific diseases.

### Functional implication of hub cells on thalamic input processing

The above sections have demonstrated that hub attacks are significantly more effective in disrupting circuit-wide synchronization in the spontaneous synchronized case. In this section, we set to test whether this observation is general enough, and also valid for the case where the synchronized activity is generated by realistic sensory input. Towards this end the we innervated the NMC circuit by 574 thalamic fibers (the thalamo-cortical, TC, input). These TC axons project mostly to neurons in lower layers 3 and 5 (**Fig. 4A**), where some neurons might receive up to 750 thalamic synapses. Each reconstructed axon is making synapses on dendrites that are adjacent to its path (**Fig. 4B**; **Methods**), functionally impacting a vertically confined space (Amsalem et al., 2020).

**Figure 4.**
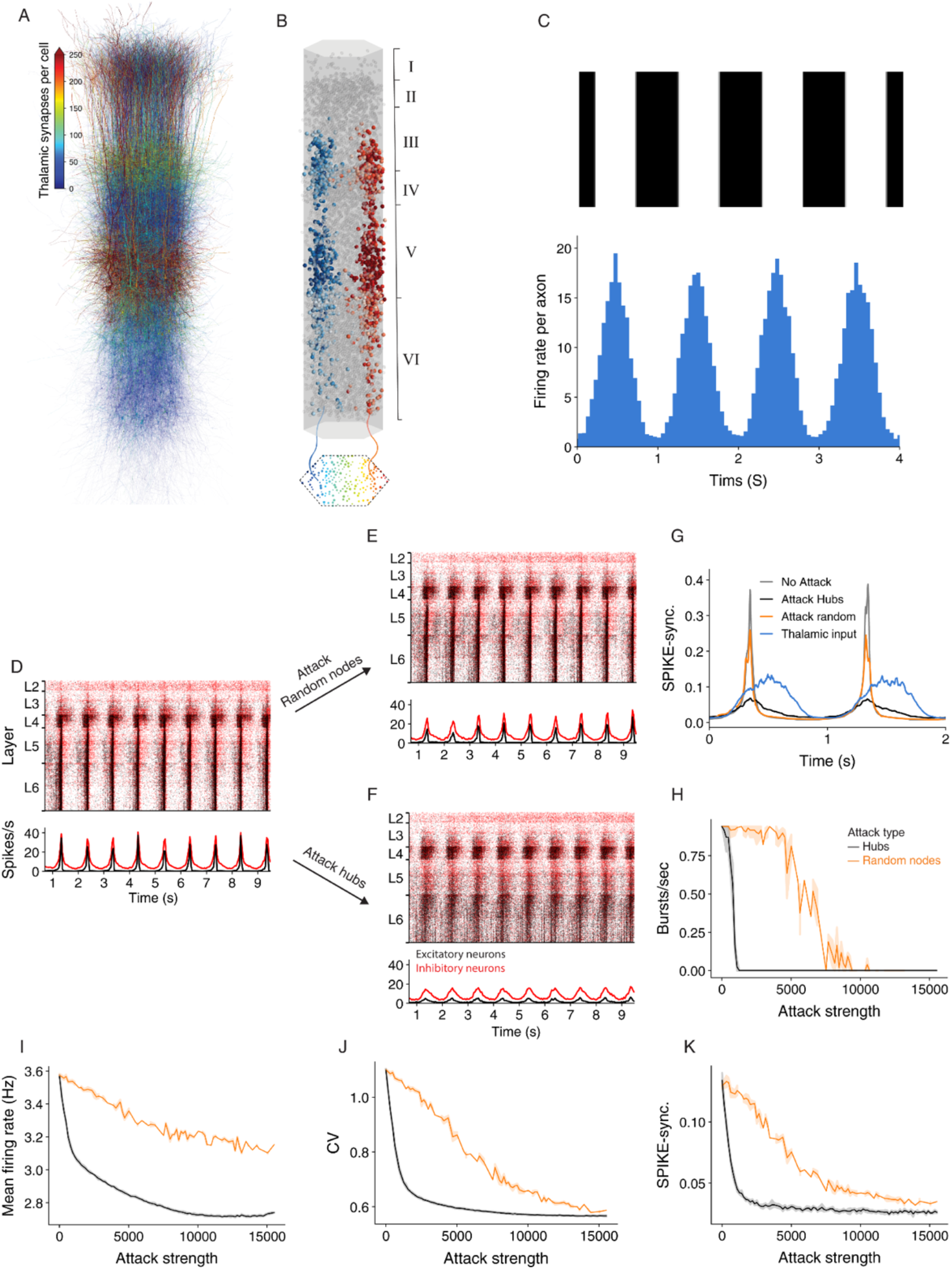
Network synchrony due to thalamic input is disrupted following attack on hub neurons. **(A)** Schematic illustration of the NMC circuit, each neuron is color-coded by the number of thalamic synapses it receives. **(B)** The spatial distribution of the thalamic input is illustrated by showing the cortical postsynaptic neurons receiving inputs from two exemplar TC axons (red and blue axons and respective colored cortical neurons). **(C)** Mean thalamic activity (bottom) simulating their response moving bars at 1 Hz (top, **Methods**). **(D)** Raster plots of the circuit responding to the thalamic input. This circuit fires asynchronously at its spontaneous state ([Ca^2+^]_0_ = 1.25 mM; see Supp. **Fig. 3A** and **Methods**). **(E-F)** The response of the circuit to the thalamic input after attack of 2,500 random nodes (E) and 2,500 hub neurons (F). **(G)** Circuit’s spike-synchronization profile (Kreuz et al., 2015) in response to thalamic input is sharpened in the intact and randomly-attacked networks as compared to that of the TC input itself (blue line) and it decreased dramatically after hub attacks (black line), even below that of the TC input. **(H-K)** Different quantifications of network activity under thalamic input for hub and random attacks (as in **Fig. 2F-G**).

For these simulations we set the calcium concentration value to 1.25 mM; this results in the network being in a spontaneous asynchronous state (As in **Supplementary Figure 3A**; see also Figure 15 and Figure S12 in Markram et al. 2015). We then simulated a grading drifting at 1Hz by generating the firing rate of the thalamic axons from an inhomogeneous Poisson process with a time-varying rate that followed a sinusoidal function (**Fig. 4C; Methods**). The circuit responded by following the oscillatory input firing with highly time-locked synchronized bursts of spikes at 1Hz (**Figure 4D**). We repeated the simulation following the attack on 2,500 random neurons (**Fig. 4E**) or 2,500 hub neurons (**Fig. 4F**). In the random attack the circuit continued to follow the oscillatory input and the response remained synchronized to the input. However, following attack on hub neurons, the circuit response was much less synchronized.

We next quantified the circuit activity using SPIKE-synchronization time profile in response to the thalamic input for different cases (**Fig. 4G**). The blue line in this figure shows the spike-synchronization measure of the TC axons whereas the grey line depicts the spike-synchronization of the cortical neurons. We found that the circuit strongly sharpens the synchronicity of the thalamic input (**Fig. 4G**, compare gray to blue line), and that attacking random nodes only slightly reduced this sharpening (orange line). In contrast, attacking hub neurons reduced the synchronization in response to the TC input dramatically (**Fig. 4G**, black line); this case is even less synchronized than the thalamic input itself (**Fig. 4G**, compare black line to blue line). We further conducted a complete set of simulations while attacking random or hubs neurons, and quantified different functional features (**Fig. 4H-K**). Hub attacks were much more destructive compared to the random attacks and caused a larger change of all measures. Interestingly removing hub neurons reduced the SPIKE-synchronization profile to a value which is lower than that of the input, showing the strong dependence of the circuit ability to follow and sharpen synchronized input on hub neurons. These results highlight the importance of hub neurons in processing sensory input, clearly demonstrating that the integrity of the cortical hub neurons is critical for the fast and reliable response of the cortical circuit to sensory information (see **Discussion)**.

## Discussion

We introduced in this work a network-based approach to investigate the relation between a cortical circuit structure to its function by removing cells according to different criteria in a highly detailed simulation. Additionally, we proposed a network-based approach for identifying potentially interesting neurons. Our analysis shows that the importance of a neuron in maintaining synchronous activity is more affected by their embedding in the network (e.g., the neurons’ in/out degree, its “hubness”) rather than strictly by their physiology.

To examine the importance of different cells and layers, we simulated the network activity after removing hub cells globally or from specific layers and compared the results to control models where random cells were removed. We discovered that hub neuron attacks have the largest change of structural network measures, leading to loss of the Small-World property of the simulated NMC (Figure **1C-E**). Accordingly, attacks on this population resulted in the largest decrease in the network synchrony, firing rate and number of bursts (Figure **2**). The attack changed the network response from spontaneously synchronous to asynchronous, resulting with no bursts and reduced CV (Nolte et al., 2019).

Among attacks targeting specific layers, mimicking a more biologically plausible scenario, we found that attacking L5 hub-neurons resulted in the largest effect on all functional measures (Figure **3D-G**) and therefore is the most distributive attack. We believe that this phenomenon is rooted in the high interconnectivity of L5 excitatory neurons (Figure **3A**). Interestingly the specific genetic profile of L5 excitatory neurons is widely used to optogenetically target and record from this subset of neurons (de Vries et al., 2020), and open the possibility of examining our predictions by specifically silencing this subpopulation while recording from the neocortex.

Hub neurons play an important role not only in maintaining spontaneous network oscillatory activity but also in the processing of sensory input from other brain areas. When thalamic (sensory) drifting sinusoidal input impinged on our modeled cortical circuit, the circuit not only followed the thalamic synchrony, but resulted in activity that was more synchronized. We then found that attacking hub cells caused a significant reduction of the synchrony of the cortical column with respect to the oscillation of the TC input (Figure **4G-K**), eventually resulting in activity that is less synchronized than the input. Nevertheless, random attacks seem to have a minor effect on thalamic input processing by the network, demonstrating yet again the robustness of the cortical microcircuit to random cell death. These results highlight the functional role of hub neurons in fast processing of sensory information in the cortical microcircuit.

Our findings can also be seen as a demonstration of how network science theory is implemented in realistic networks. As the modeled cortical microcircuit was shown to maintain the small world property with a high clustering coefficient and short mean path (Gal et al., 2017), we now can systematically characterize how the targeting of fundamental components of a small-world network, the hubs, indeed leads to loss of this characteristic in the neural microcircuit, and results in major functional disruptions. We now have concrete evidence that hubs fulfill their theoretical key role in a highly detailed biological model. Loss of Small-World between brain regions was shown to be related to neurodegenerative diseases using fMRI data (Sporns, 2013b). Our experiments suggest that a decrease in network clustering and an increase in the mean short path can cause functional failures also at the microscale level of resolution.

To conclude, while neurons are usually characterized according to genetic markers, morphology and physiology (Berg et al., 2020; Yuste et al., 2020), we showed how a specific structural measure such as the number of synapses that defines hub cells, has a direct effect on the network functionality. As hub cells are specific subtypes of neurons (Gal et al., 2017), it is possible to use *in vivo* cell-specific knockout experiments to explore the behavioral implications of the neural network functional disruption suggested in our work.

## Methods

### NMC connectedness measures

The general connectedness of the NMC was characterized by identifying connected components of the network. A strongly connected component is a group of nodes in which any node is reachable from any other node through a directed path (a series of nodes and directed edges). Intuitively, a strongly connected component reflects a group of recurrently interlinked neurons that could give rise to an anatomical module with functional specialization.

### Small-world properties

The first property of the small-world analysis is based on the length of the shortest path *l_ij_* between pairs of nodes in the network. A path length between two nodes in the network is expressed as the number of connections along that path. To generalize this property for the entire network, the characteristic path length (*l*) of a network was used, which is the mean shortest path length averaged over all pairs of neurons

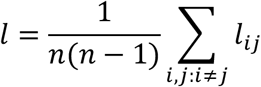

Clearly, this measure is well-defined in connected networks where any node is reachable from any other node. The NMC network initially contained a single giant component of 31,329 ± 5 neurons that were mutually reachable (99.95% ± 0.01% of all neurons; see above). The second property of the small-world analysis is captured by the tendency of nodes to cluster together. The local clustering of individual nodes measures the level at which the neighbors of a node are interconnected among themselves. Let the binary (unweighted) adjacency matrix of a directed network be denoted by *A*; then the local clustering coefficient *c_i_* of a node *i* is defined as,

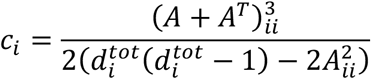

where 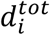 depicts the total degree (in-degree + out-degree) of node *i*. Essentially, in directed networks, this definition reflects the ratio of the number of triangles among a node and its neighbors to the number of all possible triangles that could have been formed (Fagiolo, 2007). The value of *c_i_* ranges from 0 (none of the neighbors are connected to each other) to 1 (all neighbors are mutually connected). The network-wide clustering coefficient (*c*) is computed by averaging over all local clustering coefficients

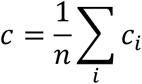

### Hubs versus the reference random attacks

To selectively attack the highly connected “hub-neurons” we started by ranking all neurons according to their total degree (total number of pre- and post-synaptic cells). Then, we performed several attacks at different strengths (the number of removed hubs). In each attack, given required number of cells to attack (*S*, attack strength), we removed the top degreed cells. To test the significance of the results observed in hub attacked networks we performed several types of random attacks for comparison. In the first, most naïve random attack, for each hub attack at strength *S* we performed 10 matching random attacks in which randomly selected *S* neurons were attacked. In the second, for each hub attack we counted the number of excitatory *S*_E_ and inhibitory *S*_I_ neurons that were attacked (*S* = *S*_*E*_ + *S*_*I*_); we then performed 10 matching random attacks in which *S*_*E*_ excitatory neurons and *S*_*I*_ inhibitory neurons were selected randomly.

### Structural analysis statistical tests

To compare the structural disruption of hub attacks to that of random node attacks (**Fig. 1**) we compared the structural metric values (mean path length and clustering coefficient) of the strongest attacks. Specifically, we took the ten strongest hub attacks (*n_1_* = 10), and the ten matching random attacks for each strength (*n_2_* = 100). For both metrics (path length or clustering coefficient) a *two-tailed Wilcoxon rank sum test* indicated that the disruption was greater for hub attacks than for matching control (*n_1_* = 10, *n_2_* = 100, *P* < 0.001).

For random edge comparisons (**Supplementary Fig. 1**) we took the 100 strongest attacks (*n_2_* = 100) and compared to the ten closest hub attacks (*n_1_* = 10). In agreement with the previous control, also here, a *two-tailed Wilcoxon rank sum test* indicated that the disruption was greater for hub attacks than for matching control (*n_1_* = 10, *n_2_* = 100, *P* < 0.001), for both metrics.

### Dense model of neocortical microcircuit (NMC)

Simulations were performed on a two-week-old previously published model of a neocortical microcircuit. Full details on the constructing of the circuits and its simulation methods were described in Markram et al. (2015). The microcircuit (Fig 1B) consisted of 31,346 biophysical Hodgkin-Huxley 3D reconstrued NEURON models with around 7.8 million synaptic connections forming around 36.4 million synapses. Synaptic connectivity between 55 distinct morphological types of neurons (m-types) was predicted algorithmically and constrained by experimental data (Reimann et al., 2015). The densities of ion-channels on morphologically-detailed neuron models were optimized to reproduce the behavior of different electrical neuron types (e-types) and synaptic dynamics recorded in vitro (Van Geit et al., 2016). Simulations were run on HPE SGI 8600 supercomputer (BlueBrain V) using NEURON (Carnevale and Hines, 2006) and CoreNEURON (Kumbhar et al., 2019).

### Simulation of baseline spontaneous activity

To account for the missing long-range connections and missing neuromodulators, neurons were depolarized with a noisy somatic current injection of 100% of first spike threshold (Markram et al. 2015 Figure 15). In addition, synapses spontaneous release probability was modified by setting the extracellular calcium concentration [Ca^2+^]_o_. Two conditions were tested, [Ca^2+^]_o_ of 1.25mM and 1.4mM each positioning the circuit in different activity regimes (Markram et al. 2015 Figure 15). Synaptic conductances and kinetics are as in Markram et al. (2015). Each attack was simulated twice with different randomization of the noisy step currents and timing of the spontaneous synaptic release, each time for 10 seconds.

### Simulation oscillatory thalamocortical input to the NMC

The oscillatory thalamocortical input to the NMC (Figure 4) was generated by following the same principle as in (Amsalem et al., 2016) - the spike times of the axons from inhomogeneous Poisson process with time-varying rate as,

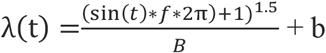

where t is time in seconds, f the frequency of the oscillatory input was set to 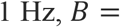 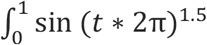 *dt* is a normalization factor, so that the mean firing rate of the oscillatory part would be equal to 7Hz, and b is the baseline spontaneous firing rate which was set to 1 Hz. For figure 5 each simulation was of 10 seconds, and conducted 5 times with different randomly generated thalamic input.

### Burst detection

We detected bursts by extracting the multivariate SPIKE-synchronization profile (Mulansky and Kreuz, 2016) for each simulation, smoothing the result using a running mean filter of ~200 ms and then counting the number of events larger than half the maximal synchronization (but at least larger than 0.15).

### Force-directed graph layout

Visualization of complex networks in an informative and meaningful way is a challenging task. How to position a large number of nodes, densely interconnected with non-obvious organization, in a two-dimensional layout that can expose inherent symmetries and structures such as hubs and clusters?

To provide a layout in which the distance between nodes (cell types) is more or less proportional to their edge weight (connection probability), we employed a Force-directed graph drawing algorithm. The algorithm is based on a physical model that assigns different forces among the nodes. On one hand, to promote attraction between connected nodes spring-like attractive forces, which depend on the distance and edge weight, are simulated. On the other hand, to avoid overlapping of nodes, repulsive forces (such as Coulomb’s law between electrically charged particles) are simulated to separate all pairs of nodes. By iteratively determining all the forces and moving the nodes accordingly, the system gets closer to an equilibrium where all forces add up to zero, and the position of the nodes stays stable. Here, we used the implementation from D3.js library (https://github.com/d3/d3-force).

### Visualization

Figures were created using Matplotlib (Hunter, 2007) . For analysis we used Python and Numpy (Harris et al., 2020)

## Acknowledgements

This work was made possible through the Patrick and Lina Drahi Foundation (PLFA), the ETH domain for the Blue Brain Project, the Gatsby Charitable Foundation, and the NIH Grant Agreement U01MH114812. We thank Noam Kahlon for the helpful discussions related to this project and for conducting the initial hub-attack experiments.

## Author’s contributions

E.G., O.A. and I.S. conceived the study and wrote the manuscript. E.G., O.A carried out the simulations and the analysis. A.S., M.L., F.S and H.M. participated in discussions and helped writing the manuscript. H.M. and F.S. developed the in silico microcircuit and provided the respective simulation framework and data.

## Supplementary Material

**Supplementary Figure 1:**
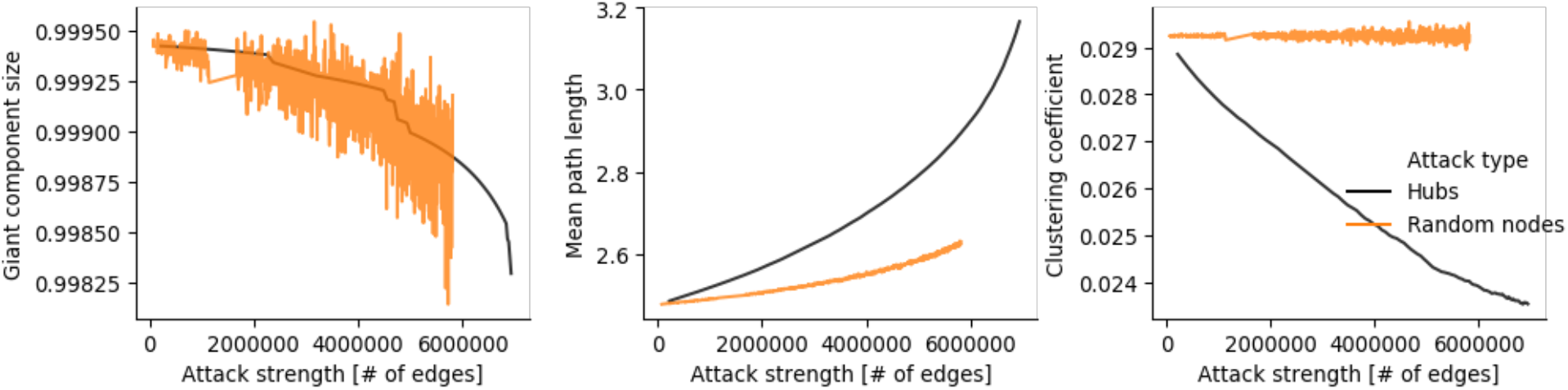
Hub attack disruption is stronger than random attacks with matching number of edges. Same as Figure 1, only the attack strength (x-axis) is measured by the number of removed edges.

**Supplementary Figure 2.**
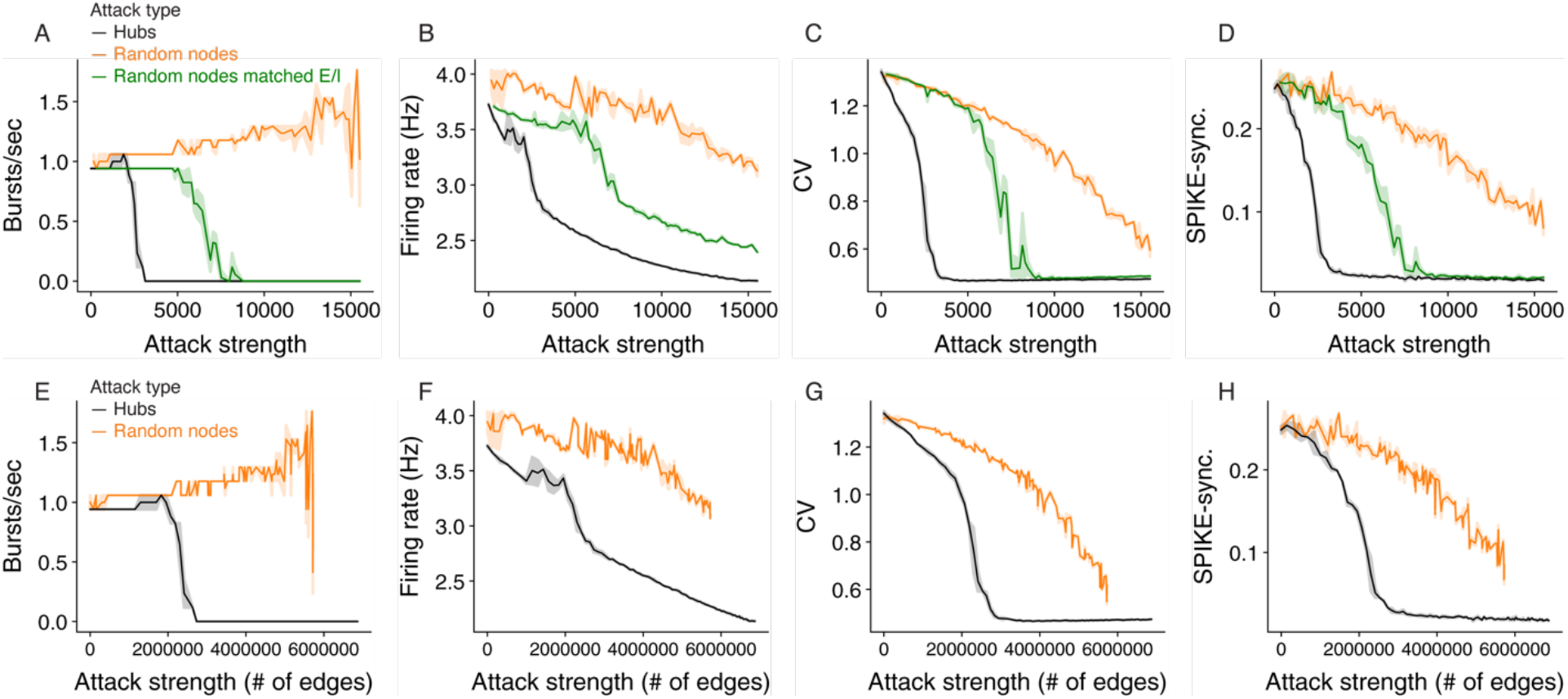
Matched E/I ratio to hub attack and matched with number of edges. **(A-D)** Hub attack (black), random attack (orange) and random attack with E/I ratio that is matched to the hub attacks (green) effects on network activity as a function of attack strength. **(A)** Average network firing rate, **(B)** The number of bursts/sec, **(C)** Correlation of variation, and **(D)** Global synchronization measure. (See **Methods** for details about the different attacks and measures). For all measures, hub attack is much more impactful. (**E-H**) same as in Figure 2, only the attack strength (x-axis) is measured by the number of removed edges.

**Supplementary Figure 3.**
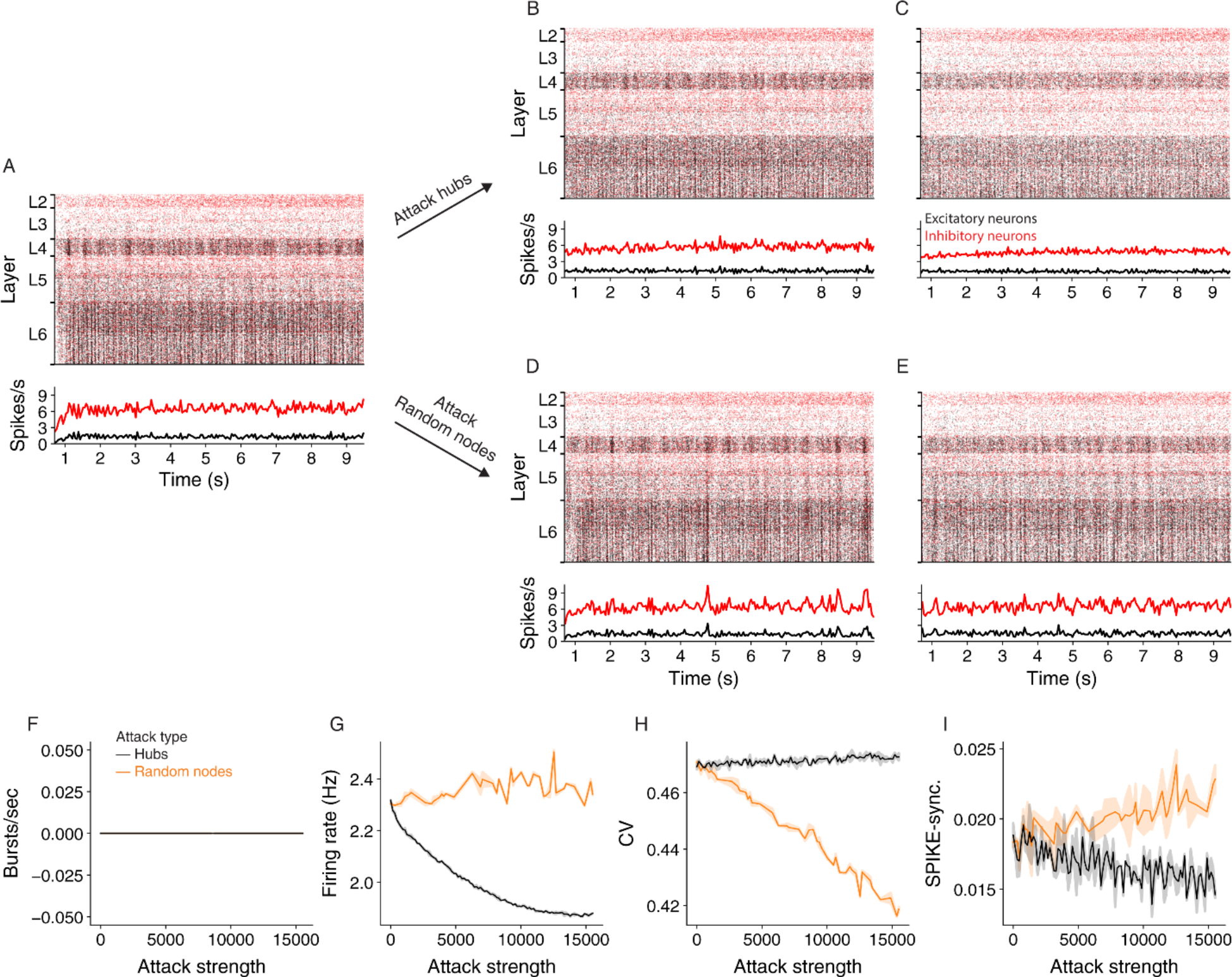
Effect of hub attacks on the dynamics of the NMC network in the a-synchronous regime. **(A)** Raster plot (top) and PSTH (bottom) of the NMC during spontaneous asynchronous state (**Methods**). **(B,C)** Same as (A) after attacking 3,134 and 8,149 hub neurons, respectively. **(D,E)** As in **(B,C)** but for respective random attacks. **(F-G)** Impact of hubs (black) versus random (orange) attacks on network activity as a function of attack strength. **(F)** On the number of bursts/sec, **(G)** On average network firing rate, **(H)** On correlation of variation **(I)** On global synchronization measure (**Methods**). For all measures, hub attack is much more impactful.

